# Epidermal microstructure and tactile sensitivity in the glabrous skin of hands and feet

**DOI:** 10.64898/2026.05.27.728083

**Authors:** James RE Hall, Gabriela Gonzalez Salas Duhne, Olivia M Wallis, Clare Howarth, Hannes P Saal

## Abstract

Glabrous skin on the human palms and soles is specialised for mechanical interaction and is characterised by papillary ridges, an absence of hair follicles, and a thick stratum corneum. While gross morphological differences, such as the increased thickness of the plantar stratum corneum, are well-documented, the fine-scale structural features associated with individual ridges and whether they help explain differences in tactile perception remain poorly understood. Here, we employed optical coherence tomography, automated image segmentation, and 3D reconstruction to generate high-resolution models of the stratum corneum and viable epidermis across six sites on the hand and foot for 27 participants. Our morphological analysis revealed that while the stratum corneum was thicker on the foot sole than the hand, the thickness of the viable epidermis remained remarkably consistent across all sites. Furthermore, papillary ridge width increased in a distal-to-proximal fashion and was larger on the foot. We also found that internal ridge architecture was considerably more pronounced than the topography observable on the skin surface. Correlating these structural parameters with psychophysical measures, specifically absolute force and two-point discrimination thresholds, demonstrated distinct drivers for tactile sensitivity. Force thresholds were primarily governed by a combination of skin thickness and innervation density, and additionally on the foot by papillary ridge width and internal ridge depth. In contrast, spatial acuity was best predicted by innervation and papillary ridge width. These findings provide a precise quantification of glabrous skin morphology and offer new insights into the process of mechanotransduction by highlighting how tissue architecture might contribute to human tactile perception.

## Introduction

The glabrous skin of the human palms and soles represents a sophisticated sensory interface, balancing the competing demands of mechanical durability and high-fidelity tactile feedback. Both regions share a common structural blueprint, characterised by a much thicker stratum corneum (Fruhstorfer et al., 2000) than in non-glabrous skin (Czekalla et al., 2019), prominent papillary ridges (Cauna and Mannan, 1961), and a lack of hair follicles. However, beyond these similarities, both skin regions serve different functional roles. Hands have evolved to serve precise manipulation and tool use (Kivell, 2015), whereas the feet are optimised for weight-bearing and postural stability (Wright et al., 2012). These differences in function likely underlie differences in skin morphology (Boyle et al., 2019) and neural innervation (Corniani and Saal, 2020), which in turn might affect sensory function.

The relationship between the skin’s architecture and the resulting perceptual experience remains an area of active investigation that can help resolve fundamental questions about which skin features help transduce tactile stimuli. Additionally, research in this area has implications for diagnosing peripheral neuropathies, designing tactile prosthetics, and advancing ergonomic tools.

Past research has mainly focused on gross structural measures, such as epidermal thickness, which can be measured relatively easily using non-invasive techniques (Lintzeri et al., 2022). Indeed, thicker skin has been linked to elevated perceptual force thresholds (Strzalkowski et al., 2015; Game et al., 2025). However, smaller structural features related to ridge morphology are harder to quantify and their potential role in sensory transduction has so far not directly been explored. Previously, modelling work has suggested that the undulating structure of the dermis/epidermis border, where low-threshold mechanoreceptors are also located, might affect the local strain distribution and thereby affect sensory function (Jobanputra et al., 2020; Maeno et al., 1998).

Other properties of the skin have also been explored in previous research. For example, harder skin has been associated with higher force thresholds (Strzalkowski et al., 2015), likely because harder skin disperses deformation due to contact over a larger area (Wynands et al., 2022). Harder skin has also been associated with worse tactile spatial acuity (Lévêque et al., 2000). Another driving factor, especially in tactile acuity, is the neural innervation of the respective skin region. Spatial acuity is highly correlated with tactile innervation density across the whole body (Corniani and Saal, 2020), and partially explains perceptual variability seen across different glabrous skin regions on the hand (Craig and Lyle, 2002). Similarly, individual differences in spatial acuity on the finger appear mostly driven by finger size, where smaller fingers lead to receptors being more densely packed in the skin (Peters et al., 2009).

Taken together, skin on the foot sole is generally thicker, harder, and less densely innervated compared to the palm of the hand, and one would therefore expect that tactile perceptual thresholds are higher on the foot sole compared to the palmar surface of the hand. However, direct comparisons between foot and hand sensitivity are rare in the literature and often do not test all potential factors mentioned above. Smaller structural features, such as those linked to individual ridge dimensions are virtually unexplored. In fact, there is a general lack of investigations into the variability of glabrous skin morphology across different sites, despite the considerable variation that is likely affecting sensory transduction.

To address these questions, our goal for the present study was two-fold: first, to characterise skin microstructure across different glabrous skin sites of the hands and feet; second, to correlate the obtained measures with two perceptual thresholds, absolute force and spatial acuity, to determine the morphological factors influencing tactile perception.

## Results

### Epidermal microstructure

We developed a method that allows visualisation and quantification of epidermal microstructure in 3D, resolving structural features of the two main epidermal layers (stratum corneum and viable epidermis), including internal features, such as limiting, intermediate, and transverse ridges up to the depth of the dermal/epidermal junction (see Figure 1A for an illustration). Our image acquisition and processing pipeline works by obtaining 2D OCT image stacks across skin patches of 4 mm × 4 mm size, pre-processing of individual slices, automated skin layer segmentation, and finally 3D point cloud reconstruction and surface meshing to generate 3D models of the stratum corneum and viable epidermis (see Figure 1B for illustration and Methods for further details). These 3D models revealed clearly visible microstructural features, mostly related to ridge morphology, across both the stratum corneum and viable epidermis (Figure 1D,E,F).

**Figure 1.**
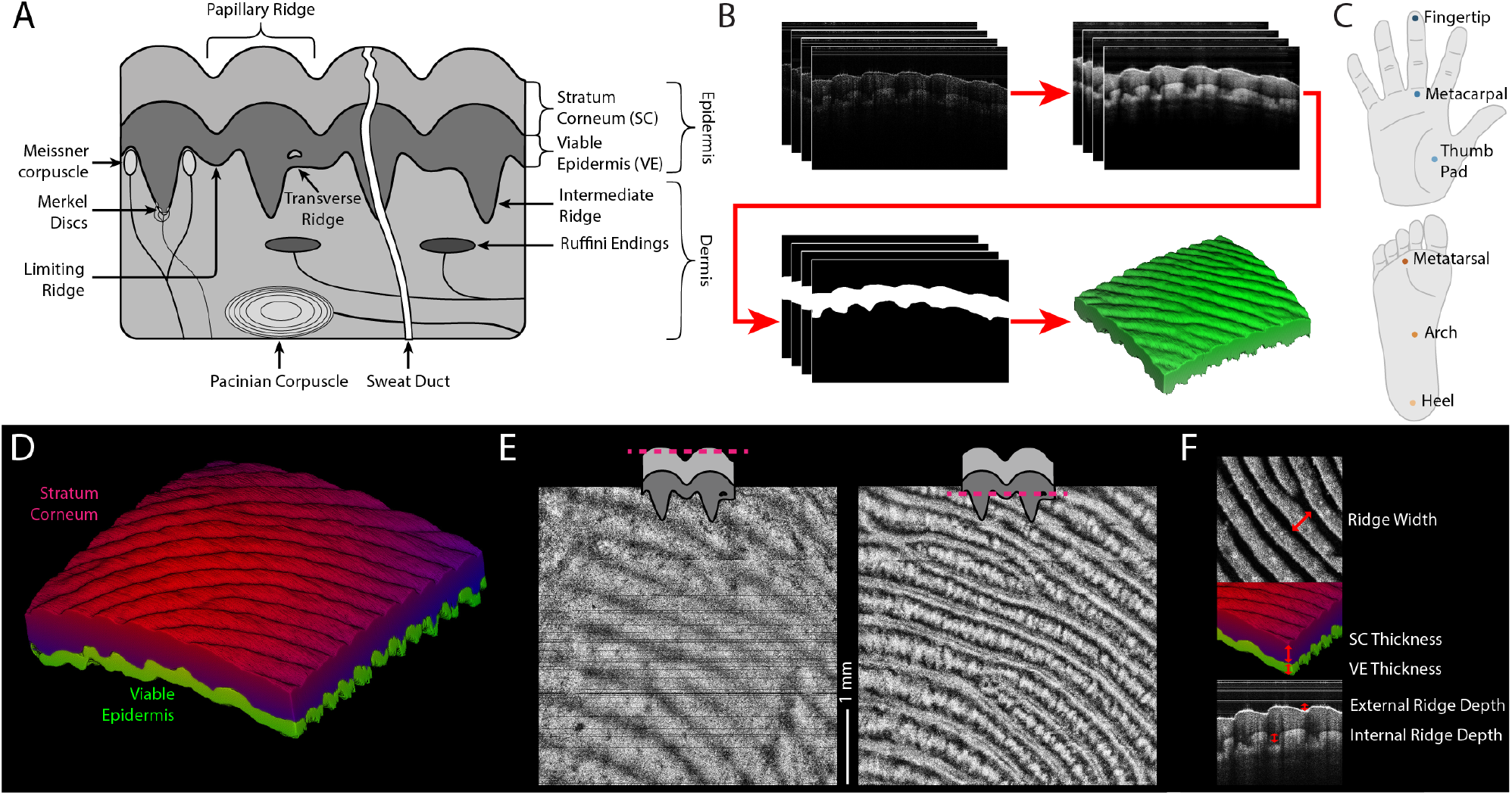
3D imaging of epidermal microstructure. **(A)** Basic structure of glabrous skin, showing different skin layers, ridge microstructures such as papillary, limiting, intermediate, and transverse ridges, as well as locations of mechanoreceptors, some of which (Meissner corpuscles and Merkel discs) are arranged with respect to the ridge structures. **(B)** Illustration of the data processing pipeline. OCT image stacks are recorded from 4 mm × 4 mm patches of skin, pre-processed to improve contrast and remove imaging artefacts, automatically segmented into skin layers (shown here: stratum corneum), and then reconstructed as full 3D models. **(C)** Six skin sites across the hand and foot were imaged: the middle fingertip, the middle metacarpal region, the thumb pad, the middle metatarsal region, the arch, and the heel. **(D)** Full 3D models allow investigation of epidermal microstructure across both the stratum corneum (red) and viable epidermis (green), here taken from fingertip of participant 9. **(E)** En-face views of skin layer boundaries are generated by slicing the OCT image stacks along layer borders determined from the 3D models. Shown are the apical surface of the stratum corneum (left) and the basal surface of the viable epidermis (right) of the fingertip for participant 13. **(F)** Quantitative measures of skin microstructure taken from OCT data. Ridge width was measured from en-face images of the stratum corneum. Stratum corneum and viable epidermis thickness was measured from the 3D models. Internal/external ridge depth measured as deflection from a smooth fit to the model apical and basal surfaces.

We used this pipeline to generate 3D models for 27 participants across six different skin sites spread across the glabrous skin: on the right hand, these were the tip of the middle finger, the middle metacarpal, and the thumb pad; on the right foot, these were the middle metatarsal, the middle arch, and the heel (see Methods for details and Figure 1C for an illustration). While these regions, as with all glabrous skin, can be characterized by the presence of papillary ridges (Figure 2A), the layout and structure of these ridges clearly varies across skin sites on the hand and foot (Figure 2C,D). For example, Figure 2C shows clear differences in the structure of the stratum corneum between the heel, which has a clear and pronounced ridged structure, and the arch, which is much flatter and thinner. Furthermore, skin microstructure in turn varies between people, even at the same location: for example Figure 2C shows deep grooves on the apical surface of the thumb pad with papillary ridges barely evident for participant 9; in contrast Figure 2D shows that participant 13 exhibits a papillary ridge structure much more similar to that of other sites.

**Figure 2.**
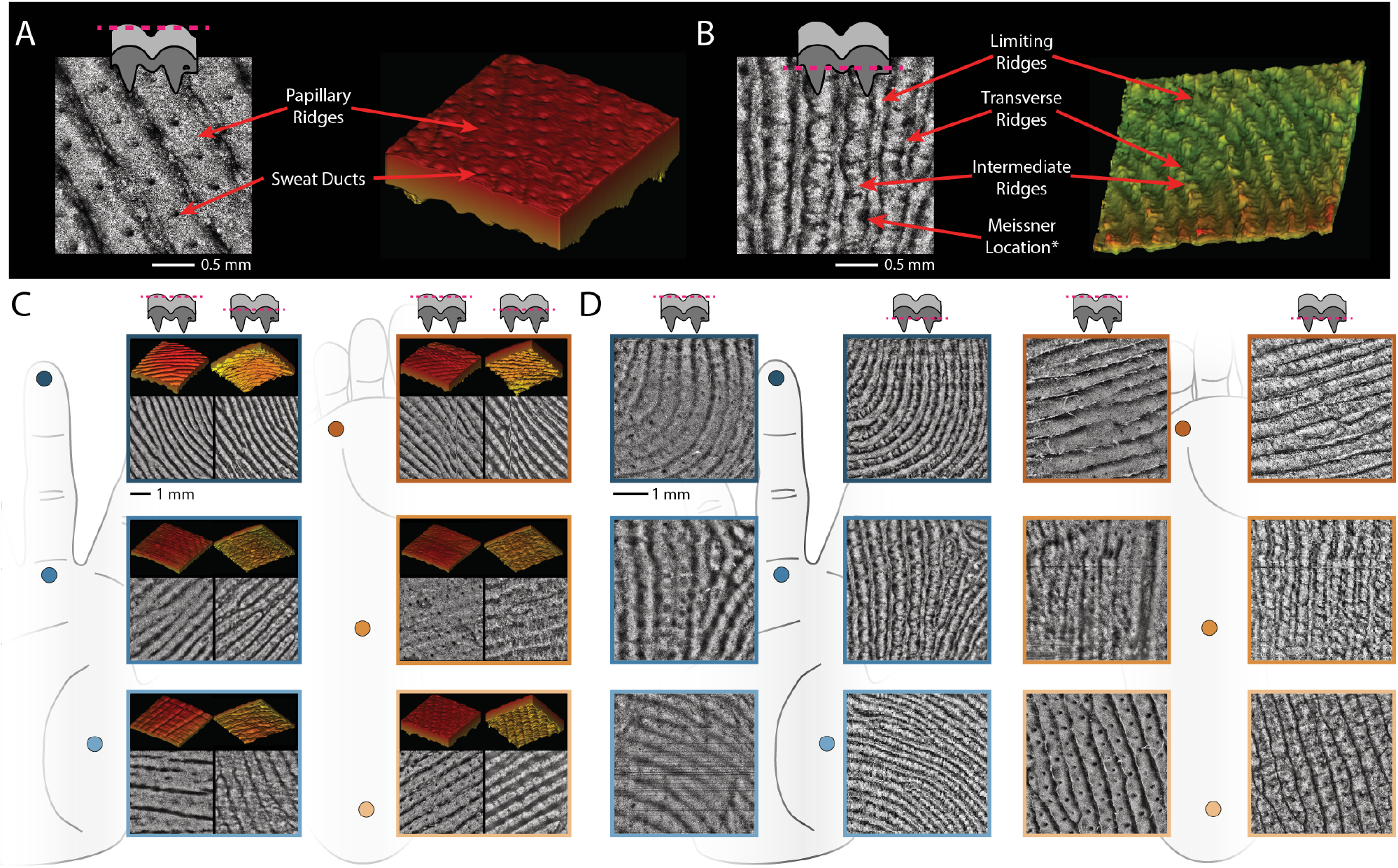
Variation in glabrous epidermal microstructure. (**A**) En face image (left) and 3D model (right) of the stratum corneum from the heel of participant 9 showing the papillary ridges of the apical surface and clearly visible holes where sweat ducts exit the epidermis. (**B**) Labelled scans of the basal viable epidermal surface of the middle metacarpal of the hand taken from participant 17. Both limiting ridges and intermediate ridges are visible, duplicating and reflecting the papillary ridge structure. Also visible are transverse ridges running orthogonal to intermediate and limiting rides. The en-face scan (left) shows distinct dark spots which are likely the locations of Meissner corpuscles. (**C**) Apical and basal en-face images (4 mm2) and 3D models of the stratum corneum of participant 9 from all sites of the hand and foot, clearly showing differences in ridge structures and thickness. (**D**) Apical surface of the stratum corneum and Basal surface of the viable epidermis (4 mm2) showing the duplication of papillary ridges to intermediate ridges at most sites and the variation in structure between the two (different participants used at each site).

The viable epidermis follows the same broad structure as the stratum corneum, however, on its basal surface (the dermal-epidermal boundary) additional ridges can be found between those seen in the stratum corneum. These so called intermediate ridges run between the limiting ridges (which reflect the valleys of papillary ridges) found on the basal surface of the viable epidermis (see illustration in Figure 1A and example in Figure 2B). In our data these intermediate ridges were often evident, however, there was considerable variation across skin sites and between participants. For example, we never observed clear intermediate ridges on the heel whose basal surface followed the apical surface (see heel example in Figure 2D, see further text in Discussion). Finally, we often found transverse ridges connecting between the limiting and intermediate ridges; these transverse ridges likely provide structural support orthogonal to the direction of the papillary ridges. Between these ridges sometimes candidate spots for Meissner corpuscles appear in the data as dark spots (similar to spots shown in Infante et al., 2023).

We calculated five structural measures from the 3D models (Figure 1F): thickness of both the stratum corneum and the viable epidermis, calculated as the average layer thickness across the full skin sample; papillary ridge width, measured from manually traced papillary ridge patterns of the stratum corneum and then calculating the distances between them; and finally external and internal ridge depth, calculated as the average deflection of the skin from a smooth curve at the apical (external) or basal (internal) surface of the stratum corneum. We also measured the hardness of the skin at all imaged sites using a durometer, as a measure of the mechanical response of the skin.

#### Hardness

On average, the skin on the foot was more than twice as hard as that of the hand (*z* = −16.94; *p* < 0.001, linear mixed effects model, Wald z-test Holm corrected; see Figure 3A). On both the hand and foot a distal-to-proximal gradient was evident, with skin becoming harder the more proximal the location. On the hand, skin on the fingertip was significantly softer compared to the metacarpal (*z* = −3.98; *p* < 0.001) and the thumb pad (*z* = 4.32; *p* < 0.001), while on the foot the heel was significantly harder than both the metatarsal (*z* = − 6.98; *p* < 0.001) and the arch (*z* = −7.82; *p* < 0.001).

**Figure 3.**
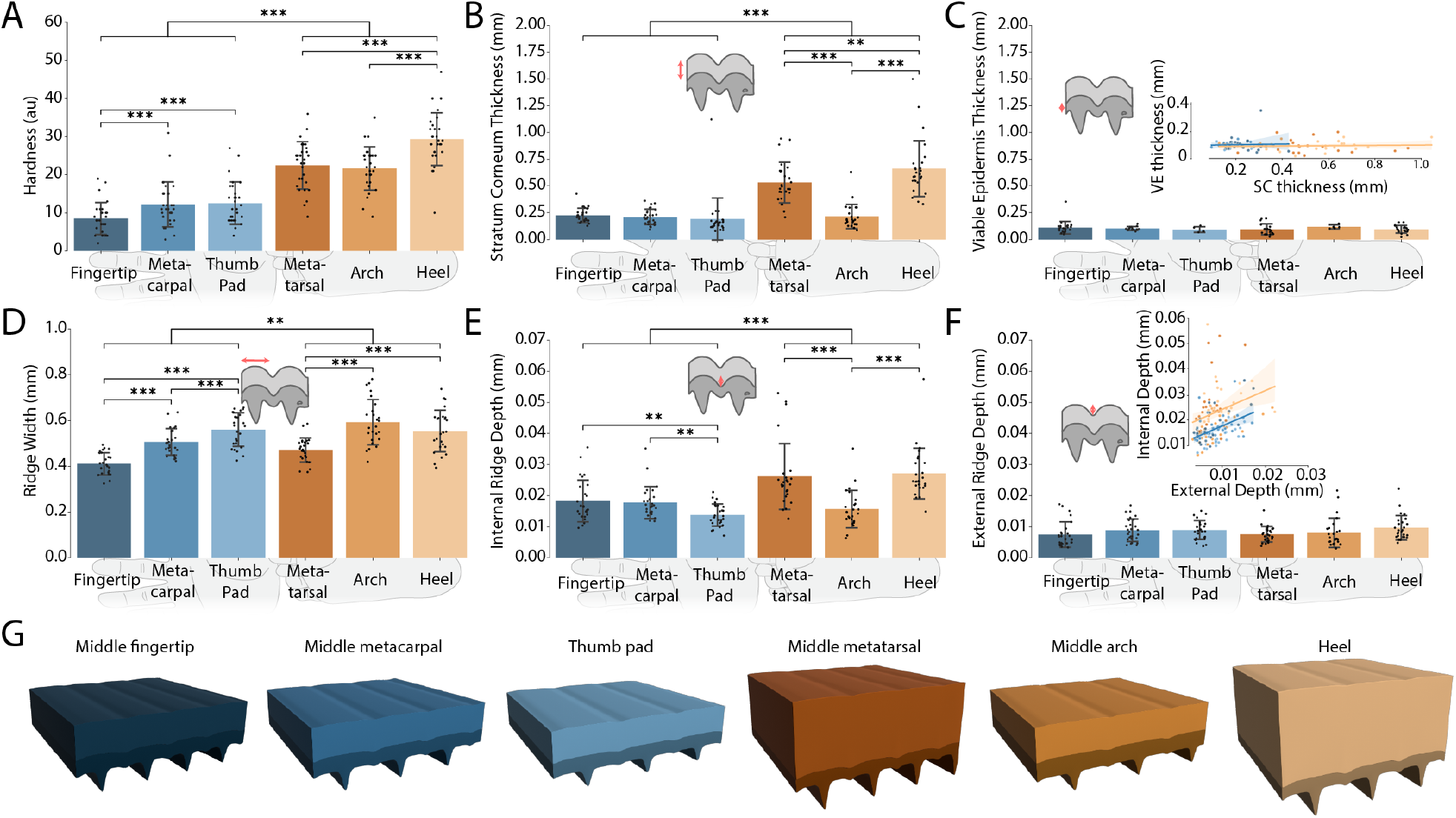
Mechanical and microstructural quantification of glabrous skin. (**A**) Skin hardness across all six sites (*n* = 174). Bars indicate averages (mean) per site, while error bars show the standard deviation. Horizontal bars at the top denote significant differences between the hand and foot, and within each body region (*: *p* < 0.05, **: *p* < 0.005, ***: *p* < 0.001). Skin on the foot is harder than on the hand. (**B**) Thickness of the stratum corneum across all sites (*n* = 156). The stratum corneum on the foot is generally thicker than that of the hand. (**C**) Thickness of the viable epidermis across all sites (*n* = 88). Note the lower number of data points as the viable epidermis could not be segmented accurately in all obtained samples. No significant differences were found. Inset: Scatter plot and trend lines (blue: hand, orange: foot) showing relationship between stratum corneum and viable epidermis thickness. Both measures are uncorrelated and the viable epidermis is generally thinner than the stratum corneum. (**D**) Papillary ridge width across all sites (*n* = 155). Ridge width increased proximally and is larger on the foot. (**E**) Internal ridge depth measured on the basal surface of the stratum corneum across all sites (*n* = 156). Internal ridges are deeper on the foot. (**F**) External ridge depth measured on the apical surface of the stratum corneum across all sites (*n* = 156). No significant differences were found. Inset: Scatter plot and trend lines (blue: hand, orange: foot) showing relationship between external and internal ridge depth. Both are moderately correlated. (**G**) Parametrically generated 3D skin layers models that accurately reflect average epidermal microstructure in terms of layer thickness and ridge depth. See Methods for details.

#### Skin layer thickness

The thickness of the epidermis was measured independently for the stratum corneum and the viable epidermis. The stratum corneum was significantly thicker on the foot and showed greater variability: while there were no significant differences across the hand with all areas displaying a uniform thickness of on average 210 µm, all three tested areas on the foot sole differed significantly from each other (all *p* < 0.01), with the heel the thickest, followed by the metatarsals and the arch being of relatively similar thickness to the hand (Figure 3B). In contrast, the viable epidermis appeared to be of a constant thickness of around 107 µm at all tested locations (Figure 3C), with no significant differences between the hand and the foot (*z* = 0.78; *p* = 0.434), or within any of these areas (all *p* > 0.346). As a consequence, the thickness of the stratum corneum and the viable epidermis were uncorrelated (*r* = − 0.04, see inset in Figure 3C). This suggests that the differences in epidermal thickness reported across these sites in the literature are due to differences in the thickness of the stratum corneum alone. Indeed, adding the measured thickness values of both the stratum corneum and the viable epidermis, results in epidermal thickness estimates closely matching those reported in the literature using a variety of different methods (Lintzeri et al., 2022). Finally, while epidermal thickness was therefore constant across the hand, we had earlier reported differences in hardness across these sites, suggesting that skin hardness cannot be explained by differences in thickness alone.

#### Ridge width

The width of the papillary ridges varied significantly between the hand and foot (*z* = −3.16; *p* < 0.01) as well as across most tested sites within a region (all *p* < 0.001), with the exception of the heel and arch which had similar ridge widths (*z* = 1.83; *p* = 0.067). We observed a broad distal-to-proximal gradient on both body parts, with average ridge width increasing from 411 µm on the fingertip and 471 µm on the metatarsal, respectively, to 560 µm on the thumb pad and 554 µm on the heel Figure 3D). Papillary ridges were on average 45 µm wider on the foot compared to the hand.

**Table 1.** Perceptual thresholds and microstructural measures across skin sites on the hand and foot. Average values (*±* standard deviation) of all measures across the hand and foot, averaged over participants.

| | Force Threshold (g) | Two-point Threshold (mm) | Hardness | SC Thickness (mm) | VE Thickness (mm) | Ridge Width (mm) | External Ridge Depth (μm) | Internal Ridge Depth (μm) |
| --- | --- | --- | --- | --- | --- | --- | --- | --- |
| **Hand (overall)** | **$0.04 \pm 0.06$** | **$5.26 \pm 2.38$** | **$11.08 \pm 5.53$** | **$0.21 \pm 0.13$** | **$0.11 \pm 0.04$** | **$0.49 \pm 0.09$** | **$8.48 \pm 3.61$** | **$16.65 \pm 5.60$** |
| Middle Fingertip | $0.04 \pm 0.07$ | $2.52 \pm 0.63$ | $8.52 \pm 4.21$ | $0.23 \pm 0.07$ | $0.11 \pm 0.06$ | $0.41 \pm 0.05$ | $7.53 \pm 4.08$ | $18.33 \pm 6.71$ |
| Middle Metacarpal | $0.05 \pm 0.07$ | $6.64 \pm 1.33$ | $12.21 \pm 5.92$ | $0.21 \pm 0.07$ | $0.10 \pm 0.02$ | $0.51 \pm 0.06$ | $8.92 \pm 3.62$ | $17.80 \pm 5.13$ |
| Thumb Pad | $0.03 \pm 0.02$ | $6.62 \pm 1.90$ | $12.52 \pm 5.57$ | $0.19 \pm 0.20$ | $0.09 \pm 0.03$ | $0.56 \pm 0.07$ | $8.98 \pm 3.02$ | $13.79 \pm 3.57$ |
| **Foot (overall)** | **$0.36 \pm 0.35$** | **$12.79 \pm 2.44$** | **$24.51 \pm 7.12$** | **$0.47 \pm 0.27$** | **$0.10 \pm 0.04$** | **$0.54 \pm 0.10$** | **$8.49 \pm 3.85$** | **$23.12 \pm 9.86$** |
| Medial Metatarsal | $0.28 \pm 0.32$ | $11.96 \pm 1.59$ | $22.48 \pm 6.23$ | $0.53 \pm 0.19$ | $0.10 \pm 0.05$ | $0.47 \pm 0.05$ | $7.63 \pm 2.46$ | $26.20 \pm 10.61$ |
| Middle Arch | $0.28 \pm 0.28$ | $14.36 \pm 2.58$ | $21.66 \pm 5.66$ | $0.22 \pm 0.11$ | $0.12 \pm 0.02$ | $0.59 \pm 0.10$ | $8.11 \pm 4.70$ | $15.71 \pm 6.04$ |
| Heel | $0.50 \pm 0.41$ | $12.06 \pm 2.31$ | $29.38 \pm 6.90$ | $0.66 \pm 0.26$ | $0.10 \pm 0.04$ | $0.55 \pm 0.09$ | $9.71 \pm 3.93$ | $27.16 \pm 8.14$ |

#### Ridge depth

The depth of the papillary ridges was measured on the surface of the skin (external ridge depth, Figure 3F) as well as at the boundary between the stratum corneum and the viable epidermis (internal ridge depth, Figure 3E). We found that the external ridge depth varied little across all the sites we tested (all *p* > 0.1) with an average depth of 8.5 µm across both the hand and foot. On the contrary, the internal ridge depth varied significantly between the hand and foot (*z* = −5.10; *p* < 0.001). Across the hand the fingertip and metacarpal had very similar ridge depths whereas the thumb pad was significantly less deep (*z* = −3.50; *p* < 0.01 and *z* = −3.14; *p* < 0.01 respectively). On the foot, the heel and the metatarsal were both similar with the arch being significantly less deep (*z* = −4.89; *p* < 0.001 and *z* = 4.52; *p* < 0.001 respectively). External and internal ridge depth were only moderately correlated (*r* = 0.47 and *r* = 0.29, for the hand and foot respectively, see inset in Figure 3F).

While the thumb pad and arch yielded similar structural measures, we noted general differences in the structure of these two sites. Skin on the arch varied more extensively between participants and was smoother than other sites on the foot, often similar to hairy skin in its externally visible structure. In contrast, skin on the thumb pad showed deep grooves on the apical surface of the stratum corneum and papillary ridges were often hard to discern at all (Figure 2C), though papillary ridges appeared clearly in lower layers. The papillary ridges generally ran orthogonal to the grooves evident on the surface.

#### Parametric skin layer models

Using the microstructural measures reported above, we generated parametric epidermal skin layer models for all six sites using the average values per skin site, accurately reflecting differences in layer thickness, ridge width and depth (Figure 3G, see Methods for details). These models better capture ridge structure than idealised textbook illustrations and can support mechanical modelling efforts to further study the effect of skin microstructure on mechanotransduction.

### Predicting tactile sensitivity from skin structure

We measured tactile sensitivity for the six skin sites across the same set of participants to establish two complementary perceptual thresholds. First, we measured absolute force thresholds using von Frey monofilaments, which allow application of a specific force onto the skin. Second, we measured spatial acuity based on a two point discrimination task. We then fit linear mixed effects models to predict each threshold measure based on the structural features described above (see Methods for more details on psychophysical protocols and model fitting).

#### Absolute force thresholds

Force thresholds varied significantly between the hand and foot (Figure 4A), with the foot exhibiting force thresholds many times higher than those on the hand (*z* = −9.75; *p* < 0.001). We did not find significant differences between force thresholds within the hand (all *p* > 0.086), although it should be noted that this might be due to the low resolution of von Frey filaments (Bove, 2006) and not necessarily reflect a lack of difference in sensitivity (see Discussion). On the foot the force thresholds were identical between the metatarsal and the arch with large variability across participants. In contrast, thresholds on the heel were significantly higher than on both the other sites on the foot (*p* < 0.001).

**Figure 4.**
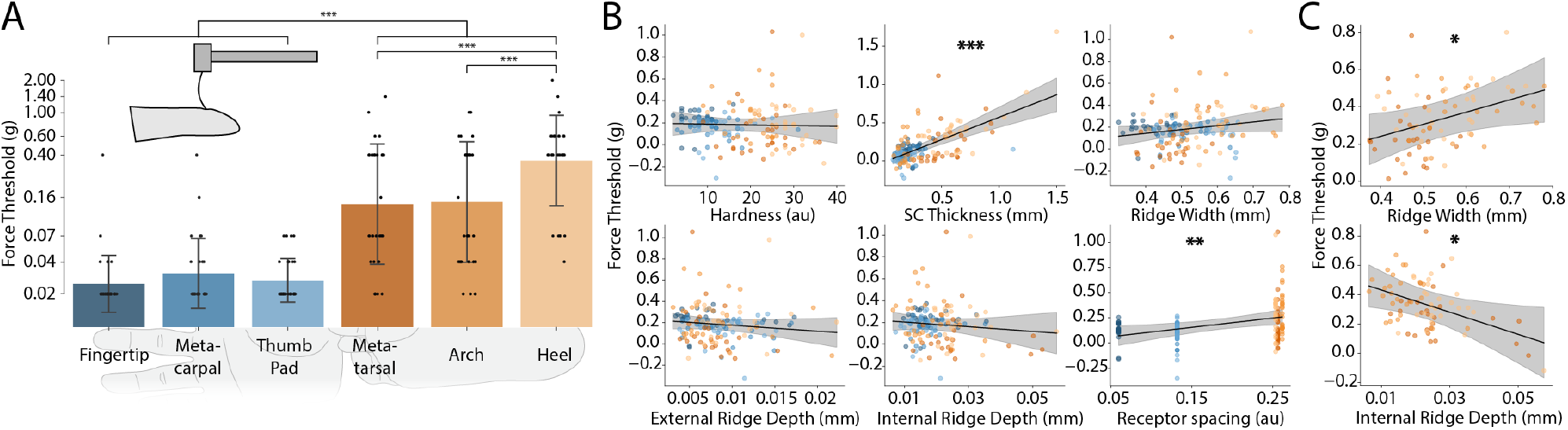
Predicting absolute force thresholds from skin structure. (**A**) Force thresholds across the hand and foot measured with von Frey filaments (*n* = 174). Coloured bars denote mean threshold and error bars denote standard deviation. Note logarithmic scale on the vertical axis to increase visibility at the low end. Horizontal bars at the top denote significant differences (***: *p* < 0.001). Force thresholds are much higher on the foot compared to the hand. (**B**) Marginal effects plots generated from a linear mixed effects model. Shown are model fits for each individual predictor (solid lines and shaded confidence intervals), after fitting all other predictors. Colours of the data points as in panel A, showing residuals for that predictor. Asterisks at the top denote significant coefficients (**: *p* < 0.005, ***: *p* < 0.001). (**C**) Significant (*: *p* < 0.05) marginal effects plots from linear mixed effects model trained on data from the foot only.

To investigate to what extent skin microstructure and other peripheral factors can account for these findings, we fitted a linear mixed effects model with six predictors (see Methods for details): skin hardness, thickness of the stratum corneum, papillary ridge width, external ridge depth, internal ridge depth, and peripheral receptor spacing. This final measure reflects tactile innervation densities expressed as the average spacing between receptors (see Methods), which have been found to strongly correlate with perceptual thresholds in previous research (Corniani and Saal, 2020; Craig and Lyle, 2002).

We found that the model predicted force thresholds extremely well (*R*2 = 60.9%) with both stratum corneum thickness (*β* = 0.59, *SE* = 0.09, *z* = 6.22, *p* < 0.001) and receptor spacing (*β* = 0.93, *SE* = 0.34, *z* = 2.73, *p* = 0.006) contributing significantly (see Figure 4B for marginal effects plots). Specifically, a difference of 100 µm in the stratum corneum thickness reflected a change in force thresholds of 0.06 g and a change in receptor spacing of 100 µm (corresponding to a change in innervation density of 100 receptors/mm2) reflected a change in force thresholds of 0.09 g.

For force sensitivity we often measured the same threshold across the entire hand (average of 0.03 g) due to the lack of resolution in von Frey filaments. Indeed, a linear mixed effects model fitted on data exclusively from the hand did not identify any significant predictors of tactile sensitivity. In contrast, a model which only considered the foot predicted local differences in force sensitivity extremely well (*R*2 = 79.0%). The stratum corneum thickness was still the main driving factor of sensitivity (*β* = 0.76, *SE* = 0.17, *z* = 4.48, *p* < 0.001). However, in contrast to the global model, receptor spacing was not significant, likely because innervation density barely varied across the different areas tested on the foot, in contrast to the big divergence between innervation of the hand and the foot (Corniani and Saal, 2020). Two additional factors were significant when only considering the foot (Figure 4C): ridge width (*β* = 0.66, *SE* = 0.30, *z* = 2.18, *p* = 0.03) and internal ridge depth (*β* = −7.56, *SE* = 3.25, *z* = −2.32, *p* = 0.02). Both affected force thresholds moderately: an increase of 100 µm in ridge width increased force thresholds by 0.03 g, and an increase of 10 µm in internal ridge depth increased force thresholds by 0.02 g. We conclude that force thresholds are mainly driven by stratum corneum thickness, with both innervation and ridge microstructure likely contributing to perception.

#### Two-point discrimination thresholds

Spatial two-point discrimination thresholds varied widely across the tested sites Figure 5A, but were fairly consistent between participants. Similarly to force thresholds, spatial thresholds were significantly lower on the hand, compared to the foot (*z* = −20.81; *p* < 0.001). On the hand, the fingertip was the most sensitive site, able to detect gaps as small as 2.52 mm between stimuli on average. In comparison, the metacarpal and thumb pad were significantly less sensitive (*z* = 12.11, *p* < 0.001 and *z* = 12.05, *p* < 0.001, respectively), with thresholds of more than 6 mm each. On the foot, the metatarsal and heel were both significantly more sensitive than the arch (*z* = −4.37, *p* < 0.001 and *z* = 4.18, *p* < 0.001, respectively). Across all skin sites, force and spatial discrimination thresholds were moderately correlated (*r* = 0.47); this relationship appeared to be driven predominantly by the foot being less sensitive for both measures, while each individual body region displayed different patterns across the two measures.

**Figure 5.**
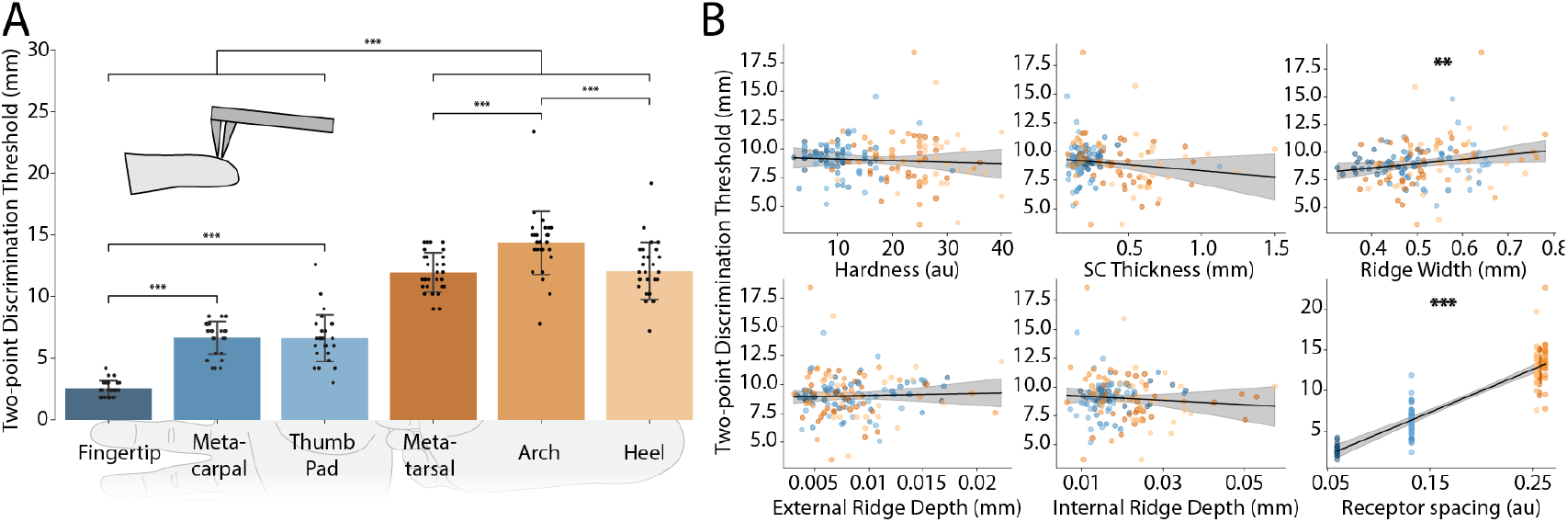
Predicting two-point discrimination thresholds from skin structure. (**A**) Two-point discrimination thresholds across the hand and foot (*n* = 174). Coloured bars denote mean threshold and error bars denote standard deviation. Horizontal bars at the top denote significant differences (***: *p* < 0.001). Thresholds are much higher on the foot compared to the hand. (**B**) Marginal effects plots generated from a linear mixed effects model. Shown are model fits for each individual predictor (solid lines and shaded confidence intervals), after fitting all other predictors. Colours of the data points as in panel A, showing residuals for that predictor. Asterisks at the top denote significant coefficients (**: *p* < 0.005, ***: *p* < 0.001).

We asked whether skin structure affects spatial discrimination by fitting a linear mixed effects model. We found that structural measures predicted spatial discrimination even more accurately than force thresholds (*R*2 = 81.7%). Across all sites, we found that thickness, unlike force sensitivity, was no longer important (*β* = −1.07, *SE* = 0.87, *z* = −1.23, *p* = 0.22). However, innervation distance was still significant (*β* = 52.95, *SE* = 3.12, *z* = 16.97, *p* < 0.001) and ridge width was also significant (*β* = 4.02, *SE* = 1.81, *z* = 2.22, *p* < 0.05). Specifically, a difference of 100 µm in receptor spacing showed a change in spatial acuity of 5.29 mm and a change of 100 µm in ridge width caused a change of 0.40 mm in spatial acuity.

In summary, both measures of tactile sensitivity are driven by neural innervation of the skin, but also by both the macro and microstructure of the epidermis. In particular, ridge width played a role in predicting both force and spatial thresholds, and internal ridge depth might be important on the foot, where the internal ridge geometry is more pronounced.

## Discussion

In this study we investigated the effects of mechanical factors on our sensitivity to touch, in particular how the microstructure of ridges in the epidermis of the glabrous skin of the hand and foot affects force and spatial sensitivity. We found that the microstructure of the epidermis varied greatly across the hand and foot and between participants, importantly we were able to define the structure of ridges across glabrous skin regions. We clearly find significant differences between these structures across all sites. Combining these structural measures and measures of sensitivity we were able to show that the microstructure of glabrous skin impacts our sensitivity to touch above and beyond that which can be explained by innervation and thickness of the skin.

### Microstructural variation across skin sites

We quantitatively characterized both macro and microstructure of the epidermis across glabrous skin in the hand and foot. Building on previous comparative work (Boyle et al., 2019), we found considerable variation across all sites tested. Hardness and overall epidermal thickness have been well studied previously (Strzalkowski et al., 2015; Vega-Bermudez and Johnson, 2004; Li and Gerling, 2023; Game et al., 2025) and our results agree with these efforts. Hardness was much higher on the foot compared to the hand, with some variation across sites. While previous studies had measured the overall thickness of the epidermis and noted thicker skin on the foot (Lintzeri et al., 2022), here we measured the thickness of the stratum corneum and viable epidermis separately. Interestingly, the viable epidermis appeared to have the same thickness across all sites while the thickness of the stratum corneum differed multiple-fold, suggesting that the main driver of epidermal thickness is the stratum corneum. Notably, the stratum corneum was thickest in areas of the foot that experience high forces during gait (Cleland et al., 2023), an observation that appears to extend to the whole body (Maiti et al., 2019). This supports the idea that the stratum corneum acts as the main protective layer to outside forces. While hairy skin exhibits a much thinner stratum corneum compared to glabrous skin, the viable epidermis appears of similar thickness in hairy skin (Robertson and Rees, 2010) as estimated here, suggesting that it might be relatively uniform across the body.

We also quantified microstructural features of the papillary ridges themselves, such as their width and the depth of both their external and internal geometry. Most of these microstructural features displayed considerable variation across both the hand and the foot. Ridge width varied across almost all the sites tested, increased along the distal-proximal axis, and was larger on the foot. The distal-to-proximal gradient has previously been described on the hand (Cummins et al., 1941; Gutiérrez-Redomero and Alonso-Rodríguez, 2013); wider ridges on the foot have been observed (Galton, 1892), but not extensively characterised. In contrast, the depth of the ridges on the skin’s outer surface, characterised as external ridge depth here and sometimes referred to as surface roughness in other studies, showed little variation across skin sites at 8.5 µm on average. Our estimates lie in the middle of previously estimated values, some lower (Maiti et al., 2019) and others higher (Derler and Gerhardt, 2012), with discrepancies likely due to different methodologies, size differences in the skin patches, or participant demographics. Notably, ridge depth appears to increase with age at least on hairy skin (Li et al., 2006). In any case, it appears clear that internal ridge geometry, measured here as the depth of the ridges on the boundary between the stratum corneum and the viable epidermis, is generally much more pronounced than the external geometry. Ridges were especially deep in the heel, which also displayed the most well defined ridges of any site, potentially as a result of the hardness and thickness of the skin there. Interestingly, ridge depth varied in a manner consistent with habitual mechanical loading across body sites. On the hand, the fingertip and metacarpal, sites frequently involved in object manipulation (Gonzalez et al., 2014), exhibited the deepest ridges, while on the foot, the heel and metatarsal, which experience regular loading during gait and balance (Cleland et al., 2023), showed a similar pattern. Deeper ridges might prevent delamination of skin layers by increasing inter-digitation (Moreno-Flores et al., 2024), which might be especially important in skin regions typically undergoing high forces. We also note that the peak-to-peak undulation of the stratum corneum at its basal surface can reach 60% or higher of the total thickness of the viable epidermis directly beneath it (see Figure 3G), causing very strong inter-digitation of this layer with the dermis below, even without taking into account additional microstructural features such as the intermediate ridges.

Apart from these easily quantifiable microstructural features, we also made a number of qualitative microstructural observations. In the stratum corneum, sweat ducts were easily visible in the OCT images, appearing as bright corkscrew-shaped structures, but this layer was otherwise devoid of other noticeable features. The viable epidermis, on the other hand, showed extensive microstructural variation, both between different skin sites and between participants. First, intermediate ridges were visible in many, but interestingly not all successful segmentation models. When they were visible, the depth of intermediate ridges was generally not as pronounced as in typical depictions derived from histology (see Figure 1A and Cauna and Mannan, 1961). Since some of our models and their associated raw data showed no visible intermediate ridges at all, it is possible that intermediate ridges extended too deep to be resolved clearly by OCT. Alternatively, it has been shown that intermediate ridges differ in their keratin expression patterns from other ridges (Swensson et al., 1998), which might interfere with their visibility in OCT images. Finally, it is possible that there is more diversity in skin microstructure than is generally evident in the selected 2D slices shown in the literature: some histology studies show more complex structure often without clearly identifiable intermediate ridges (Boyle et al., 2019; Wang et al., 2011). Some of this structure might be explained by transverse ridges, which connect intermediate and limiting ridges. These are not identifiable in 2D slices, which typically run orthogonal to the papillary ridge orientation, but have been clearly observed in 3D scanning electron microscopy volumes (Nagashima and Tsuchida, 2011; Nagashima et al., 2011).

Transverse ridges were often identifiable in our 3D models of the viable epidermis, resulting in a lattice-like structure of the dermal-epidermal junction. It has been suggested that the structure of transverse ridges differs across skin sites (Nagashima and Tsuchida, 2011; Nagashima et al., 2011), a promising avenue for future investigations into detailed three-dimensional skin structure. Finally, we observed dark spots in the OCT data, consistent in their position and size with the location of Meissner corpuscles (Infante et al., 2023).

### Tactile sensitivity of glabrous skin

By measuring both force thresholds and spatial acuity, we aimed to form a broad picture of sensitivity across glabrous skin. Indeed, force and two-point discrimination thresholds were only moderately correlated, implying that these measures tap into different aspects of tactile sensing and processing. For example, while the heel showed the least spatial acuity, it was better at force detection than the arch. Nevertheless, both measures showed clear differences between the hand and the foot, with the hand much more sensitive.

For absolute force thresholds, we did not find significant differences across the hand. Partly, this result reflects the fact that young people, our participant demographic, tended to display extremely low thresholds on the hand (Bowden and McNulty, 2013), often at the limits of resolution of the monofilaments. Other studies have reported regional differences on the hand (Giner et al., 2026), though these are still much smaller than any thresholds we recorded on the foot sole.

For two-point discrimination, we found that, as expected from previous investigations (Craig and Lyle, 2002; Bowden and McNulty, 2013), the fingertip was more sensitive than the rest of the hand, which in turn showed lower thresholds than anywhere on the foot.

### Linking structure and perception

Using linear mixed effects models we found significant predictors of tactile sensitivity that together explained most of the variance in perceptual thresholds for both absolute force and two-point discrimination thresholds.

Apart from the structural and mechanical features measured earlier, we included another predictor, corresponding to peripheral tactile innervation density. This was motivated by the fact that peripheral tactile neurons are required to transduce mechanical stimuli and convey this information to the brain.

As there is currently no non-invasive technique to measure peripheral innervation density, we relied on average published estimates for the different skin regions (Corniani and Saal, 2020). We then converted the peripheral innervation density into a metric representing the average receptor distance (see Methods), which is thought to best reflect influences on perception, following previous studies (Craig and Lyle, 2002; Corniani and Saal, 2020). Indeed, we found that receptor spacing was a significant predictor of both force and two-point discrimination thresholds. Its effect size on force thresholds was moderate, and indeed skin thickness explained much more of the effect. However, receptor spacing was an excellent predictor of two-point discrimination thresholds, as might be expected from the fact that inputs from multiple neurons would be needed to distinguish two points from a single contact point.

Thickness of the stratum corneum, which determines overall epidermal thickness as we have seen above, did not predict two-point discrimination thresholds, but was an excellent predictor of absolute force thresholds, as had previously been reported for the foot alone (Strzalkowski et al., 2015; Game et al., 2025).

We also tested a number of microstructural measures associated with individual ridge geometry. Ridge width might be assumed to be strongly linked to neural innervation because mechanoreceptors are located relative to geometrical features associated with papillary ridges (Infante et al., 2023). However, we found that ridge width was only moderately correlated with receptor spacing (*r* = 0.37), similar to previous efforts Dillon et al. (2001). Indeed, differences in measured ridge width between the hand and the foot were much smaller than estimated differences in innervation density (Corniani and Saal, 2020). Our analysis therefore tested whether ridge width predicted perceptual thresholds above and beyond any effect explained by innervation density alone. For force threshold we found no effect overall, but did find a significant moderate contribution of ridge width on the foot (Figure 4). For two-point discrimination, ridge width was the only significant predictor other than receptor spacing, though with a small effect size (Figure 5). Previous work has demonstrated that papillary ridges may limit spatial resolution by constraining tissue movement within individual papillary ridges (Jarocka et al., 2021; Corniani et al., 2025). A similar mechanism may underlie the effects observed here, however, further experimental evidence is required to verify this.

External ridge depth was not a significant predictor in any of the fitted models; while this parameter might affect the effective contact area and the resulting friction when a surface is moved across the skin, it does not appear to govern relevant mechanics that affect force sensation or spatial acuity. In contrast, internal ridge depth was a significant predictor of tactile force thresholds on the foot, with larger undulations lowering force thresholds. This is in agreement with prior modelling work suggesting that the magnitude of undulation at the dermal-epidermal border focus strain signals (Jobanputra et al., 2020; Maeno et al., 1998), which might lead to lower thresholds. Similarly, such an effect has been proposed as being relevant in edge detection (Gerling, 2010). It therefore seems that the deeper in the skin, and therefore the closer to the relevant receptors, the more important this ridged structure becomes.

Finally, although hardness varied considerably across the hand and foot, it was not a significant predictor for any of the perceptual thresholds, presumably because other predictors explained the observed perceptual effects better. Thus, skin hardness might not be well suited to explain broad differences in perceptual thresholds across skin sites, which demonstrate a range of other structural differences as well as we have seen, but it might still affect local changes in perception when the hardness of any specific skin site changes, as demonstrated previously (Lévêque et al., 2000; Wynands et al., 2022). Alternatively, it has been argued that typical skin hardness measures, such as the one employed here, provide a measure of the bulk hardness of the skin, rather than of the epidermal layers specifically (Chatzistergos et al., 2022), with the latter being potentially much more relevant for tactile thresholds.

### Limitations

While OCT has many benefits, such as its ability to take invivo measurements very quickly and with no safety concerns, this method also comes with limitations. The number of successfully segmented models for the viable epidermis in the present study was much lower than the number of models for the stratum corneum. The main reason for this was the fact that the dermal-epidermal junction was often obscured and not clearly visible. The reason for this was not generally obvious, but it was not related to the thickness of the stratum corneum. It is possible that skin hydration is an important factor, and some studies have recommended clearing agents such as vaseline or coconut oil (Banerjee and Poddar, 2022; Welzel et al., 2004). However, in pilot testing, these had little to no effect on our data quality, so we did not employ them in this study.

While both psychophysical tasks used in this study provided robust data that demonstrated clear differences in sensitivity between different skin regions, they also showed inherent limitations. As mentioned previously, von Frey monofilaments lack resolution at lower force levels, and therefore could not accurately differentiate between the most sensitive skin sites on the hand. Thus, while we did not find differences in force thresholds across the hand, it is possible that a more sensitive method could establish more subtle differences. Furthermore, two point discrimination tasks that ask participants to distinguish between one or two felt touches have been criticised for not exclusively assessing spatial acuity where other cues might be available and for sometimes yielding inconsistent results (Lundborg and Rosén, 2004). We decided to use the classic two-point paradigm, as it allows for quick assessments and was mostly easily adapted to different skin sites, which proved difficult with alternative methods we explored.

### Future work

Having shown that epidermal microstructure strongly predicts tactile sensitivity, future work should focus on the micromechanics of ridge deformation and how these impact receptor activation to arrive at a mechanistic explanation. Detailed mechanical modelling should prove useful in this endeavour and we hope that our parametric models of skin layer geometry will help with this goal. Ultimately, combining detailed mechanics with neural activation models (see e.g. Saal et al., 2017; Katic et al., 2023) might allow for a detailed understanding of the full mechanotransduction pathway.

Future work could also consider the contribution of further biomechanical, rather than purely structural, skin properties on tactile perception. Beyond skin hardness, skin hydration and (visco-)elasticity are amongst potentially important factors (Ní Annaidh et al., 2012; John et al., 2023; Saal et al., 2025). It has also been demonstrated that different postures affect both the structure of the skin (Maiti et al., 2019) and tactile sensitivity (Smith et al., 2021; French et al., 2022). Given the extensive skin stretch present across the body (Rupani et al., 2025), future work might consider how movement constantly modulates tactile perception.

Our study focused on a relatively homogeneous participant pool, consisting of young healthy adults. However, tactile thresholds dramatically change with age (e.g. Bowden and McNulty, 2013), likely in part due to mechanical and structural changes in the skin itself (McIntyre et al., 2021; Deflorio et al., 2022). Teasing out the specific contributions of different microstructural features to the decrease in sensitivity would shed further light on the process of mechanotransduction and how it can break down. Our choice to restrict our investigation to young participants was deliberate, because other factors will likely affect tactile sensitivity, such as declines in receptor numbers (McIntyre et al., 2021; Corniani and Saal, 2020) and general sensory processing (Humes et al., 2013) that we were unable to measure.

Finally, while hairy skin shows structural differences, such as the much thinner stratum corneum and absence of papillary ridges, differences in innervation and receptor types (Corniani and Saal, 2020), and alternative mechanotransduction pathways, such as activation of tactile receptors via hair follicles (Agramunt et al., 2023), there are also commonalities that suggest shared mechanisms. For example, the dermal-epidermal boundary also exhibits undulations (Langton et al., 2017; Roig-Rosello and Rousselle, 2020; Moreno-Flores et al., 2024), potentially serving a similar function as we found in glabrous skin.

## Methods

### Participants

27 healthy participants (19 female and 8 male) with no history of skin conditions took part in this study. The participants had an average age of 21.1 years (range: 18-26) and all but 2 participants were right-handed. All provided their written informed consent to participate in the study, and for the imaging and perceptual data to be made public. The study protocol was approved by the ethical review committee of the School of Psychology at the University of Sheffield (protocol number: 058482).

### Data acquisition

All experiments took place in the Skin Barrier Research Facility, operated by Sheffield Dermatology Research at the Royal Hallamshire Hospital in Sheffield, UK. The laboratory was environmentally controlled and all tests were undertaken at 20°C room temperature and 45% relative humidity.

For each participant, sensitivity testing and imaging was completed on 6 sites, with 3 on the hand and 3 on the foot (see Figure 1C), with the full protocol taking just under an hour. These sites were chosen based on preliminary testing for varying in both structure and sensitivity. On the hand these were the centroid of the middle fingertip, the center of the thumb pad and the center of the middle metacarpal. Specifically these were defined as: the midpoint of a line extending in the proximal–distal direction from the whorl of the papillary ridges to the distal end of the fingertip, the midpoint of a line extending from the base of the thumb to the wrist crease and the midpoint of a line extending from the metacarpal of the ring finger to the metacarpal of the index finger, respectively. On the foot these were the centroid of the heel, the center of the middle arch region and the center of the middle metacarpal. The sites on the foot were defined by: the midpoint of a line extending from one side of the heel to the other in both the proximal-distal and medial-lateral directions, the midpoint of a line extending from the base of the middle toe to the bottom of the heel and the midpoint of a line extending from the tip of the middle toe to the point marked as the middle arch, respectively. Sites were marked with an ink pen prior to testing to ensure the same site was targeted in the perceptual and imaging experiments. This mark did not affect image quality.

The order of testing between body regions (hand and foot) was counter-balanced across participants and the individual site order was randomised. First, force thresholds were tested at all sites belonging to a body region, then a durometer was used to measure skin hardness, followed by two point discrimination testing, finally OCT imaging was completed on the sites. Before testing began on the initial site, the psychophysical force and two-point protocols were explained and demonstrated to the participant. Participants were blindfolded throughout the perceptual testing and asked to rest their hand palm up on a table or in the case of the foot, rest it on a footrest in a comfortable position; these positions were relaxed and maintained throughout the study to prevent the effects of posture on sensitivity, which have been previously observed on both the hand (French et al., 2022) and the foot (Smith et al., 2021).

#### Force Threshold Testing

Von Frey monofilaments (Aesthesio precision tactile sensory evaluators, DanMic Global) were used to assess force thresholds at the six sites. Participants were instructed that they would be given a “3-2-1” countdown after which a stimulus either would or would not be presented. Participants were asked to report whether they could feel the stimulus with a simple yes/no response. A practice trial was run for demonstration purposes before the start of the full protocol.

The psychophysical protocol was based on a 3-down 1-up staircase method previously proposed (Tracey et al., 2012). On the hand, the first stimulus in the sequence was 1.0 g mm^−2^, while on the foot it was 4.0 g mm^−2^. These values were used because pilot testing confirmed that these stimuli were consistently supra-threshold, while sufficiently close to the actual threshold to prevent long search sequences. At each tested stimulus level, the monofilament was presented 5 times, with the end of the monofilament being placed on the marked point for 1 second per presentation. For the first stimulus in the sequence, if at least 3 of the 5 presentations were perceived by the participant, then the following stimulus would be 3 force steps lower; otherwise the following stimulus would be increased by one force step. For later stimuli in the sequence, 3 or more out of 5 perceived touches led to a decrease of one step, and otherwise to an increase of one step. Threshold was reached when a stimulus was perceived, and the following lower stimulus was not.

#### Two Point Discrimination Testing

A two point discrimination task was used to assess spatial acuity. Analogously to force testing, participants were given a “3-2-1” countdown after which a stimulus was presented. They were asked to report whether they felt one or two distinct touch points. Touches were applied using a pair of callipers (Digital ABS AOS Caliper, Mitutoyo, UK), which were adjusted using a step size of 0.6 mm (similar to Tong et al., 2013)). The same 3-down, 1-up protocol as for force testing was employed, starting at 6.0 mm for the hand and 13.2 mm for the foot. Between each presentation the orientation of the callipers in reference to the testing site was randomised by the assessor to mitigate directional bias. The presentation of the callipers was centred on the marked point such that the middle of the two points lay on the mark.

#### Hardness

Hardness of the skin was measured using a model 1600 type OO durometer (Rex Gauge LLC, Lake Zurich, IL, USA). The durometer was placed at each skin site to give an arbitrary hardness value.

### Image Acquisition

A Vivosight Optical Coherence Tomography system (Michelson Diagnostics, Maidstone, UK) was used to image all skin sites using a 20 kHz swept-source laser at 1300 nm wavelength. 3D stacks of a 4 mm2 square region of skin were acquired at each site, composed of 800 B-scan images with 5 µm spacing and 1024 samples per A-scan. During each scan, the participant was in the same position as during sensitivity testing; hand and foot in a relaxed position resting on a table or footrest respectively. The scanner head position was secured using a tripod (Veo 3+ 263CP, Vanguard World, Christchurch, UK) and located against the skin such that the site mark was approximately in the middle of the scan. A standoff attached to the scanner head and touching the skin reduced any movement of the skin during scanning. A full 3D scan took 1 minute per skin site.

### Data analysis

#### Image processing and 3D model construction

Two Unet++ networks (Zhou et al., 2018) trained on 557 manually segmented images were used to segment the stratum corneum and viable epidermis, respectively, from each obtained 2D image slice. Before segmentation each individual image was preprocessed using custom python code; this included wavelet-FFT filtering to remove stripe artifacts common in OCT acquisitions (Byers and Matcher, 2019), histogram equalisation to increase contrast, gamma correction, and local averaging to remove speckle noise. Speckle denoising was achieved by taking the average of 4 neighbouring frames (inspired by the speckle modulation method described in Liba et al., 2017). The segmented 2D regions were then used to create full 3D volumes; any floating points were removed from the final volumes.

En-face surface images at different depths (skin surface, stratum corneum/viable epidermis border, dermal-epidermal junction) were obtained by fitting a quadratic surface to the respective skin surface and then sampling the raw 3D OCT volume data at the obtained coordinates. This process generated high quality structural images, because microstructures associated with different layers crossing the smooth sampling surface are differentially reflective in the OCT data.

### Structural measures

All structural measures were extracted from 3D segmented skin data.

Thickness of both the stratum corneum and viable epidermis was measured as the average difference in height between the top and bottom surfaces of the respective layers across the full 3D model. As each volume was 800 by 800 pixels in size, this average was calculated on 640,000 individual thickness measurements per 3D model.

Ridge depth was measured as the average absolute distance of the surface (apical or basal stratum corneum from a smooth cubic fit to the surface (specifically a 3rd degree polynomial was fit to each slice):

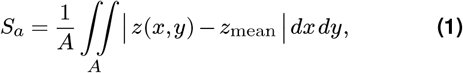

where *A* is the evaluation area, *z*(*x, y*) is the surface height function taken from the 3D model, and *z*_mean_ is the mean surface height over the evaluation area. This measure is mathematically the same as surface roughness or the arithmetical mean height (Sa, ISO 25178-2), reported in other studies. The returned value is half the peak-to-peak distance and therefore the full extent of the ridges is twice the calculated value.

Ridge width was measured as the average distance between the valleys of the ridges, with this being measured as the lowest point of each valley. Ridges were manually identified and located prior to measurement. Measurements near ridge junctions were considered outliers as they can distort valley-to-valley distances and were therefore excluded from the analysis.

#### Statistical analysis

Linear mixed effects models were fit to investigate the effects of structural skin properties on tactile sensitivity. Models were fit in Python using the *statsmodels* package (Seabold and Perktold, 2010), with the *marginaleffects* package (Arel-Bundock et al., 2024) used for marginal effects plotting. In order to investigate the effects the models were run with either two point discrimination or von Frey sensitivity as the dependant factor and ridge width, internal and external ridge depth, hardness, SC thickness, and receptor spacing as the independent variables. Receptor spacing was calculated as the square-root of the total estimated innervation density of the respective body region. On the hand, the overall density values used were: 280.9 units/mm2 for the fingertip and 57.51 units/mm2 for the palm regions (based on the values reported in Johansson and Vallbo, 1979). On the foot, the values used were: 15.25 units/mm2 for the middle metatarsal, 14.61 units/mm2 for the middle arch and 15.58 units/mm2 for the heel (based on the relative densities reported in Strzalkowski et al. (2018) scaled by the estimated total innervation reported in Corniani and Saal (2020)). Slopes were treated as fixed effects, while intercepts were modelled as random effects across participants. Post-hoc pairwise comparisons were conducted using Holm corrected Wald z-tests, using the *statsmodels* package.

#### Parametric modelling

We developed a set of equations to parametrically model the boundaries of the stratum corneum and viable epidermis, which can be adapted easily to measured dimensions from different regions. Equation 2 generates a curve, which represents the apical or basal surface of the stratum corneum and allows ridge width, ridge depth and valley shape to be adjusted in real world units:

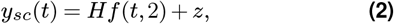

where *H* is the external or internal ridge depth to model the apical and basal surface, respectively, and *z* is the y offset from 0 and can be the offset between both surfaces and therefore the layer thickness of the stratum corneum.

The undulating shape of the ridge itself is modelled as follows:

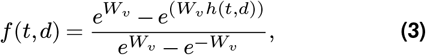

where *W*_*v*_ (> 0.1) is the valley shape constant (here set as 1.5 for all models in Figure 3), and 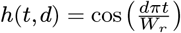, with *W*_*r*_(> 0.1) as the ridge width (valley to valley).

The basal surface of the viable epidermis doubles the ridge frequency due to the presence of intermediate ridges and needs to account for differences in depth between intermediate and limiting ridges:

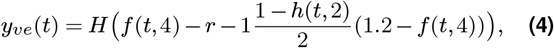

where *r*(> 1) denotes the depth of the intermediate ridges, set as a multiple of *H* (here set as *r* = 8 for all models in Figure 3).

## ACKNOWLEDGEMENTS

This work was funded by the Leverhulme Trust under Research Project Grant RPG-2022-031.

## AUTHOR CONTRIBUTIONS

Conceptualization: J.R.E.H., H.P.S.; Methodology: J.R.E.H., G.G.S.D., O.M.W., H.P.S.; Software: J.R.E.H.; Formal analysis: J.R.E.H.; Investigation: J.R.E.H, G.G.S.D., O.M.W.; Writing - Original Draft: J.R.E.H., H.P.S.; Writing – Review and Editing: G.G.S.D., O.M.W.; C.H.; Visualisation: J.R.E.H.; Supervision: C.H., H.P.S.; Funding acquisition: H.P.S.

